# Quest for the Allmitey: Potential of *Pronematus ubiquitus* (Acari: Iolinidae) as a biocontrol agent against *Tetranychus urticae* and *Tetranychus evansi* (Acari: Tetranychidae) on tomato (*Solanum lycopersicum* L.)

**DOI:** 10.1101/2021.04.08.438973

**Authors:** Viktor Van de Velde, Marcus V. A. Duarte, Alfredo Benavente, Dominiek Vangansbeke, Felix Wäckers, Patrick De Clercq

**Affiliations:** Department of Plants and Crops, Faculty of Bioscience Engineering, Ghent University, Coupure Links 653, 9000 Gent, Belgium; R&D Department, Biobest Group N.V., Westerlo, Belgium

**Keywords:** biological control, tomato, spider mites, predatory mites, greenhouse

## Abstract

The spider mites *Tetranychus evansi* Baker & Pritchard and *Tetranychus urticae* Koch (Acari: Tetranychidae) are key tomato pests worldwide. Biological control of spider mites using phytoseiid predatory mites remains challenging. The glandular trichomes on the tomato leaves and stem severely hamper the movement and establishment of the predatory mites. As a result, smaller predatory mites, able to thrive under the sticky heads of the glandular trichomes, have gained much interest. As some iolinid predatory mites were reported to feed on spider mites, we investigated the potential of *Pronematus ubiquitus* McGregor to control both *T. urticae* and *T. evansi* on tomato plants. On whole tomato plants, *P. ubiquitus* was able to suppress populations of *T. urticae*, but not of *T. evansi*. Based on the marginal number of spider mites killed in laboratory trials, the observed biocontrol effect on full tomato plants might not be due to direct predation but to a plant-mediated indirect impact. The oviposition of *T. urticae* was found to be significantly lower on tomato leaflets pre-exposed to *P. ubiquitus* as compared to non-exposed leaflets. The oviposition rate of *T. evansi* was not affected by previous exposure of the tomato host plant to *P. ubiquitus*. We demonstrated that *P. ubiquitus* reduces the population growth of *T. urticae* on tomato plants. Further large-scale field trials need to confirm the findings of the present study.

## 1. Introduction

With a worldwide production of about 180 million tons, tomatoes are among the most important vegetable crops worldwide (FAOSTAT, 2020). Tomato crops can suffer severe production losses from mite infestations, such as the broad mite *Polyphagotarsonemus latus* Banks (Acari: Tarsonemidae), several *Tetranychus* spider mite species (Acari: Tetranychidae), and the tomato russet mite *Aculops lycopersici* Massee (Acari: Eriophyidae) (Leppla et al., 2018; Navajas et al., 2012; Vervaet et al., 2021). Growers increasingly utilize integrated pest management (IPM) practices to reduce the input of chemical pesticides (Van Lenteren and Nicot, 2020). Likewise, the traditional chemical control of spider mites is shifting towards biological control within an IPM framework (Van Leeuwen et al., 2010). A cornerstone of IPM in many greenhouse crops is biological control provided by phytoseiid mites, which are the most important natural enemies used to control mite pests (Knapp et al., 2018; Van Lenteren, 2012). A plethora of phytoseiid predatory mites have been tested for their control potential against mite pests on tomato. Although pest mites like *A. lycopersici* and *Tetranychus* spp. are excellent food sources for various phytoseiids in the laboratory (Brodeur et al., 1997; de Moraes and Lima, 1983; Park et al., 2011; van Houten et al., 2013), biological control on tomato crops using phytoseiid predators is limited. The leaves and stem of the tomato plant are covered with glandular trichomes, which protect the plants from most herbivores and affect the establishment of natural enemies, including Phytoseiidae (Van Haren et al., 1987; van Houten et al., 2013). For *A. lycopersici*, on the other hand, the presence of glandular trichomes is extremely beneficial as it provides an enemy- and competitor-free space (van Houten et al., 2013).

Hence, alternative predatory mites, not impeded by dense trichomes, such as the predatory mites *Homeopronematus anconai* Baker and *Pronematus ubiquitus* McGregor (Acari: Iolinidae), have been investigated (Abou-Awad, 1979; Hessein and Perring, 1986; Kawai and Haque, 2004; Pijnakker et al., 2020; Van Lenteren and Nicot, 2020). Both predators are found naturally on tomato (Abou-Awad, 1979; Kawai and Haque, 2004) and can pre-establish in tomato crops after augmentative releases when pollen is supplemented (Hessein and Perring, 1988, 1986; Pijnakker et al., 2020). These iolinids are partly phytophagous (without causing damage) and fungivorous, enabling the mites to develop persistent populations in a crop even when prey is absent (Lorenzon et al., 2009). These characteristics provide an advantage for growers as predator populations can establish on the plant before herbivores arrive (Pijnakker et al., 2020; van Houten et al., 2019). *Pronematus ubiquitus* and *H. anconai* proved to reduce populations of *A. lycopersici* in field trials (Hessein and Perring, 1986; Pijnakker et al., 2020). *Pronematus ubiquitus* has a cosmopolitan distribution. It has has been found in North America (Acuña-Soto et al., 2017; Denmark and Porter, 1973; Mcgregor, 1932), South America (De Sousa et al., 2015; Fiaboe et al., 2007), Europe (Kumral and Çobanoğlu, 2015; van Houten et al., 2019; Vela et al., 2017), Africa (Abou-Awad et al., 1999; Ueckermann and Grout, 2007), Asia (Baradaran and Arbabi, 2009; Barbar, 2016; Gerson, 1968), and Oceania (Maynard et al., 2018). However, more research is required to confirm their potential to control the pest in greenhouse-grown tomatoes. Both *H. anconai* and *P. ubiquitus* have also been reported to feed on spider mites, although the evidence is limited. *Homeopronematus anconai* was observed to feed on eggs of the Pacific spider mite *Tetranychus pacificus* McGregor and the Willamette spider mite *Eotetranychus willamettei* Ewing (Acari: Tetranychidae) (Knop and Hoy, 1983). For *P. ubiquitus*, Dean (1957) reported feeding on newly hatched larvae of *Oligonychus pratensis* Banks (Acari: Tetranychidae). Other studies reported that a decline of spider mite populations was at least associated with the presence of predatory Iolinidae (Metwally et al., 2014; Zaher et al., 1971).

In the present study, we investigated the potential of *P. ubiquitus* to control the two-spotted spider mite *Tetranychus urticae* Koch and the red spider mite *Tetranychus evansi* Baker & Pritchard (Acari: Tetranychidae). These spider mites cause severe economic damage to tomato crops worldwide (Furtado et al., 2007; Leppla et al., 2018). The former is a cosmopolitan pest, which harms a wide variety of crops. The initial damage is visible as small pale spots, leading to white or yellow discolored patches and the leaves becoming brittle (Meck et al., 2012). *Tetranychus evansi* is an invasive pest of tropical origin specialized in solanaceous plants (Migeon et al., 2009). The damage pattern also consists of white chlorotic spots, as is the case for *T. urticae*, but without discoloration or crumbling of the leaves (Ximénez-Embún et al., 2016). Additionally, *T. evansi* produces extensive webbing, ultimately resulting in ‘mummifying’ the host plant (Ferrero et al., 2011).

The dense cover of glandular trichomes on the plant’s surface is an integral part of the tomato plant’s defense mechanisms against herbivores. Apart from causing physical obstructions for many arthropod visitors, these trichomes also release defense compounds such as volatile metabolites and proteins (Glas et al., 2012). Interestingly, *T. urticae* and *T. evansi* differentially deal with the tomato plant’s defenses. Upon attack by most strains of *T. urticae*, jasmonic acid-regulated defenses are being induced in the tomato plants, which results in a negative effect on the mite’s performance (Li et al., 2002). *Tetranychus evansi*, on the other hand, suppresses the induction of plant defenses to such an extent that the level of defense in infested plants is lower than the so-called baseline levels in control plants (Sarmento et al., 2011). *Tetranychus evansi* populations have currently only been described as suppressors of plant defenses (Alba et al., 2015; Sarmento et al., 2011), while both defense-inducing and defense-suppressing *T. urticae* strains have been reported (Kant et al., 2008). In most natural *T. urticae*-populations, however, the suppressors are not the dominant genotype (Alba et al., 2015; Kant et al., 2008).

The suppression of plant defenses also tends to be beneficial for competitors of the herbivore responsible for the suppression. These defense-inducing competitors no longer need to cope with the plant’s defensive responses. This effect is even observed among *T. urticae* populations, as inducing females have a higher oviposition rate on leaves that previously had been or are still infested with a strain that suppresses plant defenses (Alba et al., 2015; Kant et al., 2008). Moreover, it can also promote predation, as Ataide et al. (2016) showed that *Phytoseiulus longipes* Evans (Acari: Phytoseiidae) consumed more spider mite eggs when the eggs were deposited on defense-suppressed plants as opposed to plants with induced defenses (Blaazer et al., 2018). As *P. ubiquitus* also feeds on plant tissue (Brickhill, 1958; Flaherty and Hoy, 1971), it can be expected to influence the plant’s defense response. There is, however, no information available on what plant responses are being induced (or suppressed). Therefore, besides a direct predation experiment in the laboratory, we also investigated the impact of previously exposing tomato leaflets to *P. ubiquitus* on the reproduction of both *T. urticae* and *T. evansi*. In a subsequent greenhouse trial, we evaluated the overall control potential of *P. ubiquitus* on whole tomato plants.

## 2. Materials and methods

### 2.1. Rearing methods

All cultures and trials were done using tomato plants of the cultivar “Merlice” (De Ruiter, the Netherlands), grown in greenhouses of the Greenlab facilities at Biobest, Westerlo, Belgium. For *T. evansi*, the Brazilian strain “Viçosa – 1” was used, reared on tomato. The *T. urticae* strain was reared on either bean (*Phaseolus vulgaris* L.) or tomato as specified in the description of the individual trials. The spider mite colonies were maintained in the laboratory at 22 ± 1°C and a 60 ± 10% relative humidity (RH). For maintaining these colonies, new plants were added every week and plants with reduced quality due to high mite infestation were removed. A strain of *P. ubiquitus* was collected in Merelbeke, Belgium (51°00’29.0”N - 3°46’05.0”E) and subsequently mass reared at Biobest Group N.V. on pollen of *Typha angustifolia* L. (Nutrimite™). An additional population of this strain has been maintained in the Greenlab greenhouse on tomato plants for over two years.

### 2.2. Predation

In this laboratory experiment, the predation capacity of *P. ubiquitus* on the egg and chrysalis stages of *T. urticae* and *T. evansi* was assessed. Tomato plants in the fourth true leaf stage (ca. 25 cm in height) were inoculated with predatory mites using five leaf arenas (2 cm x 5 cm) per plant cut from other tomato plants with high *P. ubiquitus* populations (ca. 20 mixed mobile stages/leaf arena). The leaf arenas were evenly distributed over the tomato plants to produce a uniform inoculation. To enhance establishment, *T. angustifolia* pollen (ca. 0.01 mg per plant) was provided as supplemental food for the predators. Three days later, fifteen rectangular leaf arenas (1.5 cm x 4 cm) were cut along the main veins of these plants’ leaves for each treatment and placed per three in Petri dishes on water-saturated cotton with the abaxial side up. Four treatments were set up: *P. ubiquitus* + *T. urticae* (“Pu+Tu”), *T. urticae* alone (“Tu”), *P. ubiquitus* + *T. evansi* (“Pu+Te”), and *T. evansi* alone (“Te”). Control arenas of the same size were cut from predator-free tomato plants of the same age. The number of predators per arena was either reduced (i.e. removed from the arena) or complemented to 10 adult females (i.e. by adding female predators from the rearings on tomato plants). The latter mites were transferred using a fine brush, while closely monitoring whether they still showed normal mobility after the transfer. Hereafter, a single adult female *T. urticae* or *T. evansi*, randomly selected from the colonies reared on tomato, was added per arena using a fine brush. Over the next three days, the number of oviposited spider mite eggs was counted, as well as the number of killed eggs. Killed eggs were observed as ‘deflated’ hulls and were removed each day. The twenty Petri dishes were kept in a growth chamber under the following conditions: 23 ± 1°C, 85 ± 5% RH, and a 16:8 h (light:dark) photoperiod.

An additional experiment was carried out to assess predation of *P. ubiquitus* on spider mite chrysalis stages on tomato leaf arenas. For each spider mite species, fifteen tomato leaf arenas (1.5 cm x 1.5 cm) were cut and placed on wet cotton with the abaxial side up. Ten randomly selected chrysalis stages of the spider mites, taken from the rearings on tomato plants, were placed on each leaf arena. Single adult female *P. ubiquitus*, randomly selected from the mass rearing on pollen, were transferred to ten arenas per spider mite species, leaving five arenas as controls without *P. ubiquitus*. Two days later, the molting rate of chrysalis stages was assessed in arenas with and without the predator.

### 2.3. Indirect effects

This laboratory experiment aimed at elucidating plant reponse-related effects of *P. ubiquitus* on *T. urticae* and *T. evansi*. A total of eight tomato plants (fourth true leaf stage; ca. 25 cm in height) was used for this trial. Insect glue (Tangle-trap^®^) was applied to the petioles of two leaflets per plant, chosen from the first and second true leaves, using a fine spatula. On four plants, these leaflets were inoculated with thirty adult females of *P. ubiquitus* from the mass rearing on pollen using a fine brush, the other four plants were not inoculated (control). After inoculation, a brush tip (ca. 0.01 mg) of *T. angustifolia* pollen was applied as supplemental food. After four days, all *P. ubiquitus* stages (mobiles and eggs) were gently removed with a brush while paying attention not to damage the tomato leaf tissue. The control leaves were subjected to the same brushing treatment, thus accounting for incidental damage to the treated leaves. Per treatment, ten arenas of 1.5 cm x 1.5 cm were cut along the main vein of these leaves, and a single adult female of either *Tetranychus* species was added per arena. The *T. evansi* females had been reared on tomato, whereas those of *T. urticae* originated from a rearing on bean plants. The arenas were placed on cotton with the abaxial leaf side up. After three days, the number of *T. urticae* or *T. evansi* eggs was counted. There were two treatments per species: *P. ubiquitus*-exposed versus control (no predators), resulting in a total of four treatments. One week later, another block was carried out as a repetition in time with only five arenas per treatment and both *Tetranychus* species taken from a culture on tomato.

### 2.4. Greenhouse trial

We evaluated the potential of *P. ubiquitus* to control *T. urticae* and *T. evansi* on whole tomato plants in a greenhouse trial. Sixty tomato plants (fourth true leaf stage; ca. 20 cm in height) were divided over six treatments: no inoculation (“control”), only *P. ubiquitus* (“Pu-control”), only *T. urticae* (“Tu-control”), only *T. evansi* (“Te-control”), as well as the combinations of each spider mite species with the predator (“Tu+Pu” and “Te+Pu”). The plants were grown from seeds at the Greenlab greenhouses at Biobest NV, Westerlo, Belgium, at an average temperature of 20.2°C (18 - 27°C), 51.9% (34.0 - 70.5%) RH and with 16 h of supplementary light (high-pressure sodium lamps). Thirty of these tomato plants (treatments “Pu-control”, “Tu+Pu”, and “Te+Pu”) were inoculated with mixed stages of *P. ubiquitus* using the sawdust-pollen mixture from the mass rearing, and another thirty plants were kept as controls. This first inoculation was carried out using ca. 1 g of the pollen-sawdust mixture from the mass-rearing, aiming at an initial inoculation of 800 eggs and 250-300 mobile stages of *P. ubiquitus* per plant. An additional brush tip of pollen (ca. 0.01 mg) was also applied on the leaves of the *P. ubiquitus*-inoculated plants as supplemental food for the predators. This pollen provisioning was repeated every week (after the counts). One week after the first *P. ubiquitus* inoculation, the same thirty plants (now ca. 35 cm high) were inoculated a second time with 1 g of the pollen-sawdust mixture, containing 1190 eggs and 450 mobile stages of *P. ubiquitus* to ensure a rapid establishment of the predator. The *P. ubiquitus* populations were counted starting one week after this second inoculation using a 25x macro lens attachment to a smartphone camera (Samsung Galaxy A40), thus counting all mobile stages per spot (the visible area on the smartphone screen). The day after this first count, the forty plants in treatments “Te+Pu”, “Te-control”, “Tu+Pu”, and “Tu-control” were manually inoculated with five adult female spider mites (*T. evansi* or *T. urticae* taken from the laboratory cultures on tomato and bean, respectively) evenly distributed over the plant’s leaves. A second count of the *P. ubiquitus* populations was done one week after the first count. One week after the first spider mite inoculation, the forty spider mite-treated plants were inoculated once more with five adult females of *T. urticae* or *T. evansi* (both reared on tomato). The *P. ubiquitus* and *Tetranychus* populations were monitored using the smartphone camera method over a total of seven weeks, starting the spider mite population monitoring at the third *P. ubiquitus* count. In the first week, only twenty-one spots of the *P. ubiquitus-*inoculated plants were counted due to the small size of the plants, whereas thirty spots were counted in the subsequent weeks. The number of counted leaves was also noted so that the spot counts could be extrapolated to obtain an estimate of the population on the entire plant while accounting for the smaller leaf size at the top as compared to the middle and bottom parts of the tomato plants.

### 2.5. Statistical analysis

All data were analyzed using the R programming language (R Core Team, 2019). Normality was visually assessed using normal QQ-plots, as well as using Shapiro-Wilk normality tests. Statistical significance was noted when p-values were smaller than 0.05. The predation trial data were pooled over three days and analyzed using a Kruskal-Wallis rank sum test for both *T. evansi* and *T. urticae* treatments since there were significant deviations from normality in the data and a confined number of repetitions. For the indirect effects experiments in the laboratory, the data of the two repetitions in time were pooled. The *T. evansi* data were analyzed using Student’s two-sample unpaired t-test as no significant deviations from normality and equality of variance (Levene test) were found. The *T. urticae* data were analyzed using a Kruskal-Wallis rank sum test since the data showed significant deviations from normality. The analyses of the greenhouse trial were carried out using least squares means of linear mixed-effects models with a Kenward-Roger approximation for the degrees of freedom. The data for the *T. urticae* treatment were log-transformed and corrected with + 0.1 for zero values. For all linear models, normality of the residuals and dispersion were assessed using QQ-plots and plots of the fitted values vs the residuals, respectively.

## 3. Results

### 3.1. Predation

Table 1 demonstrates the results of the predation by *P. ubiquitus* on spider mite eggs.

**Table 1:**
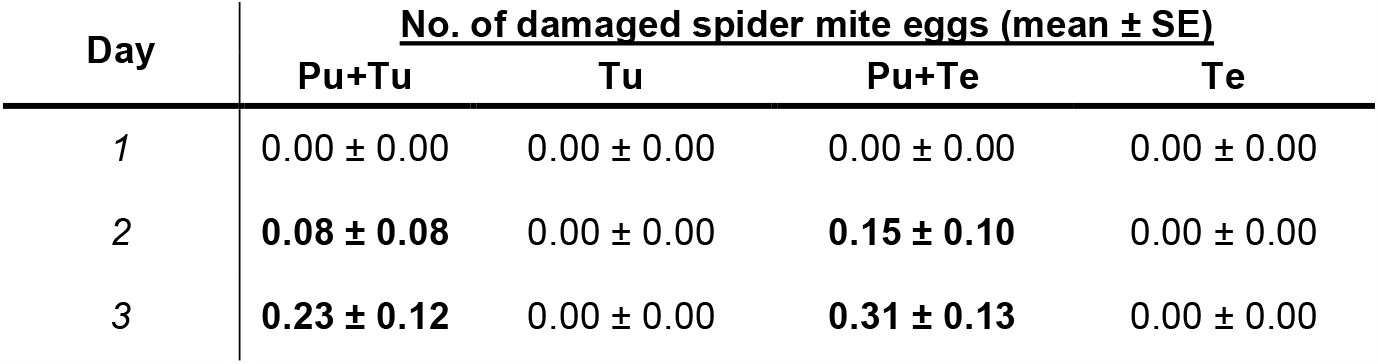
Number of damaged eggs (mean ± SE) per day of *T. urticae* and *T. evansi* when confronted with 10 *P. ubiquitus* individuals during three successive days.

| Day | No. of damaged spider mite eggs (mean ± SE) | | | |
| --- | --- | --- | --- | --- |
|  | Pu+Tu | Tu | Pu+Te | Te |
| 1 | 0.00 ± 0.00 | 0.00 ± 0.00 | 0.00 ± 0.00 | 0.00 ± 0.00 |
| 2 | **0.08 ± 0.08** | 0.00 ± 0.00 | **0.15 ± 0.10** | 0.00 ± 0.00 |
| 3 | **0.23 ± 0.12** | 0.00 ± 0.00 | **0.31 ± 0.13** | 0.00 ± 0.00 |

There were no damaged eggs in the control treatments without predatory mites. For *T. evansi*, the total number of damaged eggs over three days differed significantly between leaves colonized by *P. ubiquitus* and control leaves (Kruskal-Wallis rank-sum test, χ^²^ = 6.7308, df = 1, p = 0.009476). For *T. urticae*, the difference between the control leaves and leaves with *P. ubiquitus* was not significant (Kruskal-Wallis rank-sum test, χ^²^ = 3.25, df = 1, p = 0.07142). In this experiment, the highest recorded mean number of damaged eggs per day was 0.31 per 10 *P. ubiquitus* adult females.

### 3.2. Indirect effects

The results of the two repetitions in time done for assessing the indirect effect of predator presence on oviposition by both *Tetranychus* spp. were combined in Figure 1.

**Figure 1:**
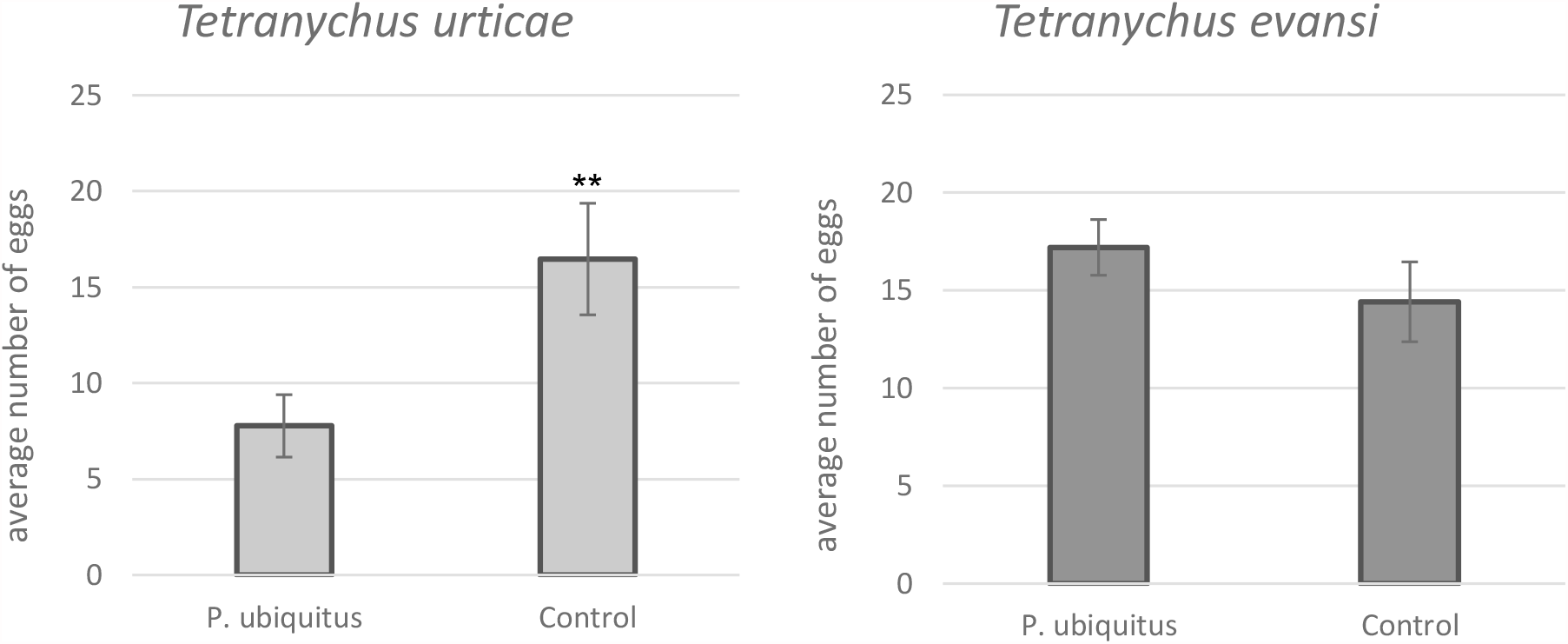
Number of eggs (mean ± SE) deposited by *T. urticae* and *T. evansi* females placed on tomato leaf arenas from plants exposed to *P. ubiquitus* vs control plants. Significant differences among treatments are indicated with ‘**’ (p < 0.05).

The mean number of oviposited *T. urticae* eggs was more than halved on the tomato leaves previously exposed to *P. ubiquitus* as compared to the control leaves without exposure to the predator. This reduction was statistically significant (Kruskal-Wallis rank-sum test, χ^²^ = 5.3658, df = 1, p = 0.02054). For *T. evansi*, these numbers did not differ (Student’s two-sample unpaired t-test, t = -1.1494, df = 25, p = 0.2613). The trial block (which also corresponded to the *T. urticae* background) did not have a significant effect (ANOVA, F = 0.0355, df = -1, p = 0.8514).

### 3.3. Greenhouse trial

The population dynamics in the greenhouse trial are shown in Figure 2 as total numbers of mites per plant over a seven week period, based on an extrapolation of the counts to the total number of leaves per plant.

**Figure 2:**
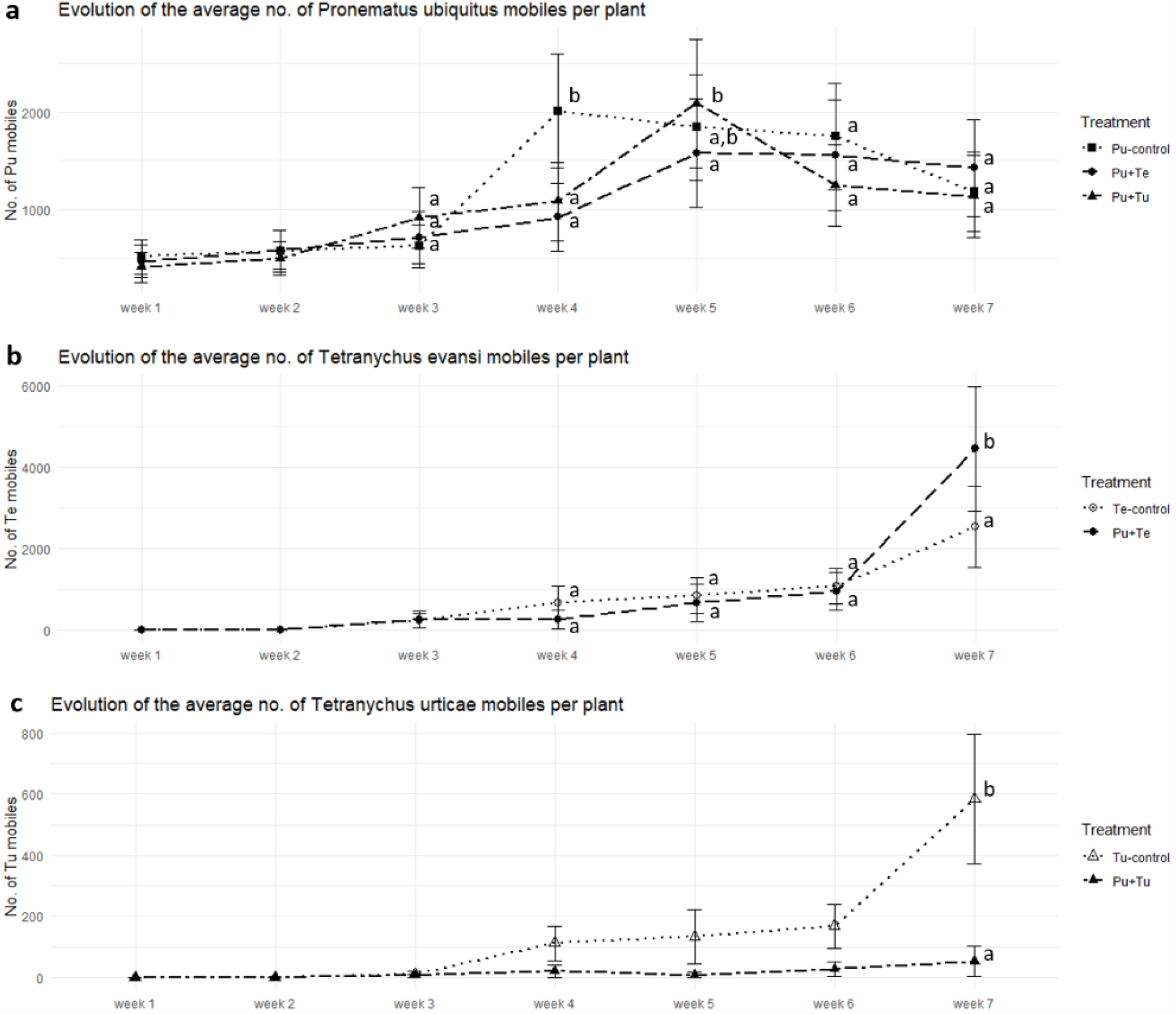
Population dynamics of *P. ubiquitus* (a), *T. evansi* (b) and *T. urticae* (c) in a greenhouse experiment on tomato. Total numbers of mobile mites per plant (mean ± SE) are shown over a period of seven weeks. Significant differences among treatments (per week in the *P. ubiquitus* and *T. evansi* graphs) are marked with different letters ‘a’ or ‘b’ (contrast after LME; p < 0.05).

Interaction between week and treatment was significant for the *P. ubiquitus* (χ^²^ = 57.845, df = 12, p < 0.001) and *T. evansi* models (χ^²^ = 21.953, df = 4, p < 0.001). In the *T. urticae* treatment, an additive model was chosen over a more complex model with interaction since the interaction was not significant (χ^²^ = 3.8561, df = 4, p = 0.4258). The *P. ubiquitus* populations followed a similar course in the different treatments, with the highest population on week 4 for the control treatment without spider mites and on week 5 for the *T. urticae*-colonized plants (Figure 2a). For *T. evansi*, there only appears to be a significant difference in numbers of mobiles in the last week, where the control plants had significantly lower numbers of spider mites than the *P. ubiquitus*-inoculated plants (Figure 2b). Plant colonization by *T. urticae* was less successful than that by *T. evansi*. It is worth noting that in the treatments with *T. urticae* one out of the ten control plants exhibited a markedly higher (i.e. a factor 20 as compared to most other plants) spider mite population. However, it was decided not to consider this replicate as an outlier as spider mite infestations in a tomato crop are known to occur in patches (French et al., 1976). The population of *T. urticae* on *P. ubiquitus*-inoculated plants was significantly lower as compared to the control plants (Figure 2c).

The trial was terminated after seven weeks of counting because at this point the *T. evansi*-treated plants had been destroyed by the spider mites.

## 4. Discussion

*Pronematus ubiquitus* has a cosmopolitan distribution on all continents (except Antarctica), including North America (Acuña-Soto et al., 2017; Denmark and Porter, 1973; Mcgregor, 1932), South America (De Sousa et al., 2015; Fiaboe et al., 2007), Europe (Kumral and Çobanoğlu, 2015; van Houten et al., 2019; Vela et al., 2017), Africa (Abou-Awad et al., 1999; Ueckermann and Grout, 2007), Asia (Baradaran and Arbabi, 2009; Barbar, 2016; Gerson, 1968), and Oceania (Maynard et al., 2018). *Pronematus ubiquitus* is able to exploit a wide range of food sources, including plant sap (Brickhill, 1958; Knop and Hoy, 1983), pollen (Flaherty and Hoy, 1971), eriophyoid mites (Abou-Awad et al., 1999; Baker, 1968; van Houten et al., 2019), and tetranychid mites (Dean, 1957). Studies have reported a negative correlation between population densities of tetranychids or tenuipalpids and *P. ubiquitus* on corn and grapes (Metwally et al., 2014; Zaher et al., 1971). However, it is unclear whether the reported correlations are associative, i.e. whether *P. ubiquitus* is the causal agent of pest mite population declines.

Our greenhouse study confirmed that *P. ubiquitus* populations could be established on tomato plants using *Typha* pollen, as reported by van Houten et al. (2019) and Pijnakker et al. (2020). Those pre-established *P. ubiquitus* populations were capable of suppressing the growth of *T. urticae* populations on tomato plants. However, based on the results of the direct predation experiments and on feeding observations during these trials, it is unlikely that this effect is solely attributable to direct spider mite killing. The mean numbers of *T. urticae* eggs killed per day by 10 *P. ubiquitus* ranged between 0.08 and 0.23 in the predation experiment. Unsuccessful feeding attempts on eggs were often observed, after which the spider mite eggs were able to hatch. A similar observation was reported by Brickhill (1958), who noted that feeding on eggs of *Tetranychus telarius* L. yielded healthy colonies of the tydeid *Lorryia ferulus* Baker. Surprisingly, however, nearly 100% of those *T. telarius* eggs hatched, which is in line with our observations. It remains to be investigated whether this feeding-but-not-killing behavior of *P. ubiquitus* on *Tetranychus* eggs affects the further development or reproduction of the spider mites. Moreover, scavenging, as well as feeding attempts on larvae and chrysalis stages were also rarely observed with the attacked prey surviving in the latter cases (Figure 3; personal observations).

**Figure 3:**
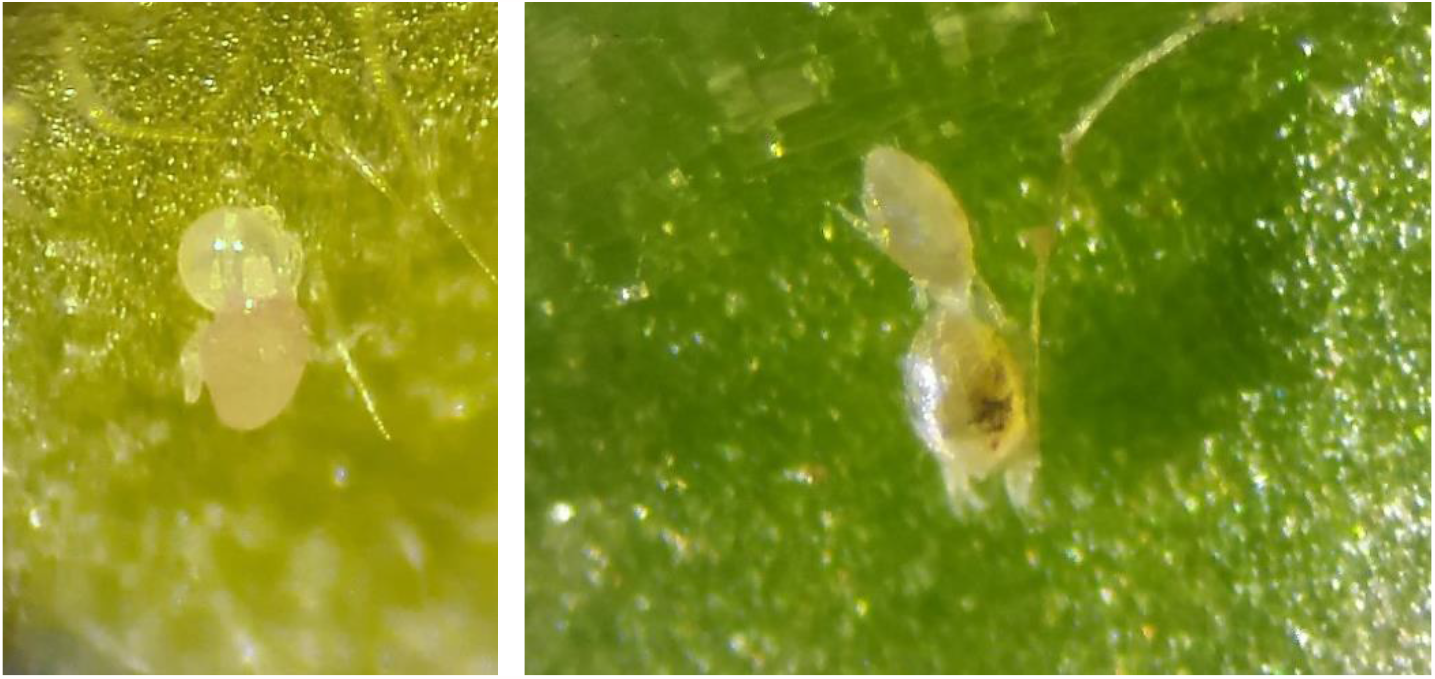
Adult female *P. ubiquitus* feeding on a *T. urticae* egg (left) and chrysalis (right).

Like many other tydeid mite species, *P. ubiquitus* is an omnivorous predator that can feed on plant sap using its stylet-like mouthparts (Brickhill, 1958; Knop and Hoy, 1983). This feeding trait is considered advantageous for the predator as it allows its populations to persist longer in the absence of prey or alternative foods. A possible effect of plant-feeding is the induction of plant defense mechanisms (Karban and Baldwin, 1997). The reproductive performance of *T. urticae* on tomato declined after pre-exposure to the plant-feeding predatory bug *Macrolophus pygmaeus* Rambur (Heteroptera: Miridae) (Pappas et al., 2015). Likewise, a lower reproduction of the western flower thrips *Frankliniella occidentalis* Pergande was reported on young tomato plants after prior exposure to another plant-feeding bug, *Orius laevigatus* Fieber (Heteroptera: Anthocoridae) (De Puysseleyr et al., 2011). A similar induced plant resistance might have been elicited in tomato leaflets exposed to *P. ubiquitus* in this study. Female *T. urticae* deposited about 50% fewer eggs on tomato leaf arenas cut from plants previously exposed to *P. ubiquitus*. The reduced growth of the *T. urticae* populations on tomato plants in the greenhouse trial may be predominantly attributable to induced responses rather than direct predation.

Further studies should address whether other key tomato pests, such as whiteflies or tomato russet mites, also experience negative non-consumptive effects from the presence of *P. ubiquitus*. We cannot exclude that the reduced oviposition by the spider mites is caused by their perception of *P. ubiquitus* cues (e.g. feces) remaining on the experimental arena. The effect of predator cues on oviposition of *T. urticae* has been observed by Ferrari & Schausberger (2013), who noted that *T. urticae* laid fewer eggs on bean leaves that contained cues of the predators *Phytoseiulus persimilis* Athias-Henriot and *Amblyseius andersoni* Chant (Acari: Phytoseiidae), as compared to leaves where predator cues were absent.

In contrast to *T. urticae*, prior exposure of the tomato host plant to *P. ubiquitus* did not affect the reproduction of *T. evansi* and its population growth was not suppressed by pre-established populations of *P. ubiquitus* on tomato in the greenhouse study. The different findings for the two spider mite species might be due to their different impact on plant defenses. Whereas *T. urticae* feeding induces defenses, *T. evansi* can suppress defenses below the plant’s constitutive levels (Sarmento et al., 2011). A laboratory study by Knapp (2007) confirms the low predatory capacity of *P. ubiquitus* against *T. evansi* since the predator was not observed to feed and did not reproduce when solely provided with *T. evansi*. There are several explanations for the observation that *T. urticae* population growth was suppressed by *P. ubiquitus*, while *T. evansi* populations were unaffected. First, the different ways the two spider mites cope with the tomato plant’s defense system would affect their population growth rate. The defense-suppressing *T. evansi* sustain their development rates by lowering the defenses of the host plant, contrary to *T. urticae* which suffer from the plant’s immune system. If *P. ubiquitus* also induces these plant defenses, *T. urticae* might suffer adverse effects, whereas *T. evansi* might not. This could explain the more expansive population growth of *T. evansi* on tomato plants colonized by *P. ubiquitus* as this downregulation of defense pathways increases the herbivore’s overall fitness (Blaazer et al., 2018; Sarmento et al., 2011). Second, *T. urticae* was observed to produce much less webbing than *T. evansi*, which facilitates the predator’s mobility. The effect of webbing on predation has been demonstrated in other systems, such as *Amblyseius swirskii* Athias-Henriot (Acari: Phytoseiidae) which is only capable of attacking spider mites that occur at the webbing’s borders (Calvo et al., 2014).

Cruz-Miralles et al. (2021) showed that the chemistry of the host plant might limit the effectiveness of an omnivorous predator as this type of predator is also expected to be susceptible to the host plant’s defense mechanisms. For instance, they noted better development of the plant-feeding phytoseiid predator *Euseius stipulatus* Athias-Henriot on a *Citrus* host plant which was more susceptible to *T. urticae* infestations, as compared to a highly resistant species (Cleopatra mandarin (*Citrus reshni* hort. ex Tan.) and sour orange (*Citrus aurantium* L.), respectively). In contrast, the rates of increase of two non-phytophagous specialist predators, *Neoseiulus californicus* McGregor and *P. persimilis*, did not differ between host plants. This research suggests that using specialist predators for biological control may be more effective when the prey feeds on better-defended plants as these plants naturally support lower herbivore densities. Direct plant defenses and protection gained from the presence of specialist predatory mites could therefore complement each other in agricultural crops. As *P. ubiquitus* is an omnivorous mite rather than a specialist predator, its effectiveness and population development on tomato plants may be limited by the strong defense responses of certain tomato varieties.

Our study indicates that pre-established *P. ubiquitus* populations can reduce the population growth of *T. urticae* on tomato plants but have no apparent effect on *T. evansi*. Preventative *P. ubiquitus* releases along with curative introductions of *P. persimilis* might be an important addition to the current spider mite IPM programs in tomato crops. Based on the results from our laboratory experiments, the effect on *T. urticae* might be primarily a result of induced defensive responses of the tomato plant rather than being due to direct predation.

## Conclusions

In conclusion, *P. ubiquitus* might play a crucial role in biological pest control on tomato, not only due to its ability to reduce tomato russet mite populations but by concurrently contributing to spider mite control. Our greenhouse trial showed that pre-established *P. ubiquitus* populations could suppress the population growth of *T. urticae* on tomato plants. Further study is required to determine whether *P. ubiquitus* can maintain *T. urticae* under economic threshold levels.

## Acknowledgments

This work was carried out alongside the Flanders Agency for Innovation and Entrepreneurship VLAIO project “BALTO”. We thank Thomas Van Leeuwen from the Department of Plant and Crops at Ghent University and Arne Janssen from the Institute for Biodiversity and Ecosystem Dynamics at the University of Amsterdam for providing us with the Brazilian *Tetranychus evansi* strain. We are grateful to Ilse Jacobs, Peggy Bogaerts, and the R&D team of Biobest Westerlo for their help and support during the trials.

